# Graph-to-Signal Transformation Based Classification of Functional Connectivity Brain Networks

**DOI:** 10.1101/541532

**Authors:** Tamanna T. K. Munia, Selin Aviyente

**Affiliations:** Department of Electrical and Computer Engineering, Michigan State University, East Lansing, MI 48824, USA

## Abstract

Complex network theory has been successful at unveiling the topology of the brain and showing alterations to the network structure due to brain disease, cognitive function and behavior. Functional connectivity networks (FCNs) represent different brain regions as the nodes and the connectivity between them as the edges of a graph. Graph theoretic measures provide a way to extract features from these networks enabling subsequent characterization and discrimination of networks across conditions. However, these measures are constrained mostly to binary networks and highly dependent on the network size. In this paper, we propose a novel graph-to-signal transform that overcomes these shortcomings to extract features from functional connectivity networks. The proposed transformation is based on classical multidimensional scaling (CMDS) theory and transforms a graph into signals such that the Euclidean distance between the nodes of the network is preserved. In this paper, we propose to use the resistance distance matrix for transforming weighted functional connectivity networks into signals. Our results illustrate how well-known network structures transform into distinct signals using the proposed graph-to-signal transformation. We then compute well-known signal features on the extracted graph signals to discriminate between FCNs constructed across different experimental conditions. Based on our results, the signals obtained from the graph-to-signal transformation allow for the characterization of functional connectivity networks, and the corresponding features are more discriminative compared to graph theoretic measures.

## Introduction

The human brain is a highly interconnected network. While early studies of neurophysiological and neuroimaging data focused on the analysis of isolated regions, i.e. univariate analysis, most of the recent work indicates that the network organization of the brain fundamentally shapes its function [1]. Thus, generating comprehensive maps of brain connectivity, also known as connectomes, and characterizing these networks has become a major goal of neuroscience [2, 3]. Complex network theory has contributed significantly to the characterization of the topology of FCNs, in particular in the assessment of functional integration and segregation [4, 5]. Specifically, graph theoretic measures such as the path length and clustering coefficient have helped to characterize small-world brain networks [6–8], and the degree distribution has been utilized to characterize scale-free networks [9]. Over the last decade, the study of FCNs through complex network theory has provided new means for discriminating between different neural dysfunctions such as epilepsy [10, 11], depression [12, 13], Alzheimer’s Disease [14, 15], and Parkinson’s Disease [16].

Although graph theoretical approaches provide an elegant way to describe the topology of functional brain networks, these measures suffer from several major shortcomings. First, most network measures are optimally suited for sparse and binary networks. Early work in the area of graph theoretic measures focused on binary networks, leading to the thresholding of networks constructed from neuroimaging studies. However, thresholding poses the problem of over-simplifying FCNs and more importantly, there is no generally accepted criterion to select the threshold [8, 17]. Moreover, the size and density of the thresholded network varies based on the chosen threshold value [4]. Recent studies show that the significance of the difference between groups is strongly dependent on the threshold parameter, i.e. the power of the statistical analysis varies with the threshold [18]. Recently, extensions of graph theoretic measures have been proposed for weighted networks to address some of these issues [19–21]. Second, it has been shown that graph theoretic measures, such as the clustering measure and the small-world parameter, are very sensitive to the size of the network, i.e. the number of nodes, and the density of the connections. Thus, comparing two networks with different edge density may lead to wrong conclusions making it difficult to disentangle experimental effects from those introduced by differences in the average degree [8, 22]. Third, graph theoretic measures are in general non-unique. An example is the small-world measure as two very different network structures may yield similar small-world parameters [23]. Finally, graph theoretic measures do not necessarily reflect the actual mechanism for flow of information in the underlying network, especially for weighted networks such as FCNs. For example, FCNs may not necessarily rely on shortest paths for communication between the nodes, and measures like the characteristic path length and the global efficiency are unable to capture this type of connectivity patterns [4, 24].

Graph-to-signal transformations have been proposed as an alternative to graph theoretic methods for network analysis [25–27]. Unlike graph theoretic measures which often result in a single number, transforming graphs into signals results in as many signals as nodes, and thus can be considered as a lossless transformation. In addition, by transforming graphs into signals it is possible to apply traditional signal processing techniques on the resulting signals in order to extract information from the networks. Both probabilistic [28] and deterministic [25, 26] methods have been proposed to transform networks into signals. Shimada *et al.* [25] and Haraguchi *et al.* [26] formulated a deterministic method based on classical multidimensional scaling, allowing the transformation of complex binary networks into time series. Under this transformation, the nodes of the network correspond to time indices in the obtained signals [25, 26, 29]. However, all of these approaches have focused on binary graphs, and have limited applicability to weighted networks that arise in neuroscience.

In this paper, we extend deterministic graph-to-signal transformations from binary to weighted networks using the resistance distance [30]. Some advantages of the resistance distance include invertibility and accounting for the global structure of the graph, thus incorporating information about multiple paths. Unlike graph theoretic measures, which map FCNs into single numbers to characterize the structure of the network, the proposed graph-to-signal transformation yields signals defined across the nodes of the network. These signals provide information about the topology of the network which can be used to extract descriptive features from the network. In this paper, we propose to implement well-known signal features such as entropy and statistical moments for these graph-signals. Finally, we apply this new transform and the accompanying features to FCNs constructed from an EEG speeded-reaction task experiment. The results indicate that the proposed graph-to-signal transformation can identify the brain regions central to error-related negativity (ERN). Furthermore, the features extracted from these signals are more discriminative compared to conventional graph theoretic measures.

## Background

### Phase Synchrony

Weighted connectivity networks were constructed from EEG data using a measure of phase synchrony. Each electrode was considered as a vertex of the graph and the weights between vertices were obtained by computing the phase synchrony between two regions. In this paper, the pairwise phase synchrony was computed by using a recently introduced time-frequency phase synchrony (TFPS) measure based on the reduced interference Rihaczek (RID-Rihaczek) time-frequency distribution [31]. For a signal *x*_*i*_(*t*), the RID-Rihaczek distribution is defined as [31]:

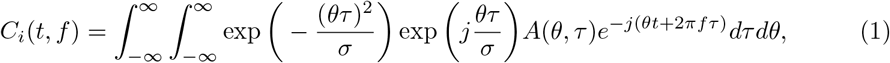

where 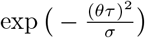 is the Choi-Williams kernel, [32], 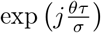 is the kernel function for the Rihaczek distribution [33] and *A*(*θ, τ*) is the ambiguity function of the given signal *x*_*i*_ and is defined as:

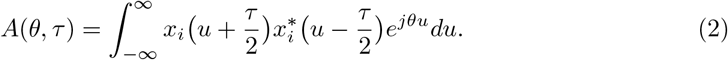

The instantaneous phase of *x*_*i*_ is computed from *C*_*i*_(*t, f*) as:

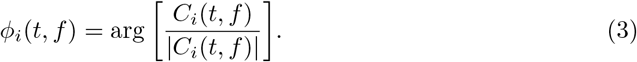

The phase difference between two signal *x*_*i*_ and *x*_*j*_ can then be computed as:

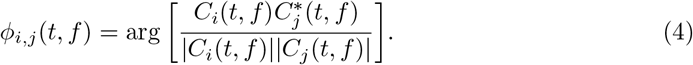

Phase Locking Value (PLV), which quantifies the phase synchrony between two signals *x*_*i*_ and *x*_*j*_, is defined as the consistency of the phase differences *ϕ*_*i,j*_ (*t, f*) across trials and can be computed as [34]:

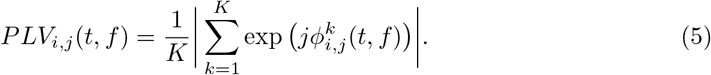

where *K* is the total number of trials and 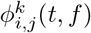 is the phase difference for the *k*th trial between 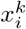 and 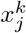 as defined by (4). Once the pairwise PLV values are computed between all pairs of electrodes, the weighted adjacency matrix corresponding to the FCN can be constructed as the average of *PLV*_*i,j*_ (*t, f*) within the time interval and frequency band of interest.

### Graph Theory

An undirected graph *G* = (*V, E*) is defined by a set of *N* nodes, *v*_*i*_ ∈ *V*, and a set of edges, *e*_*ij*_, *i, j* ∈ {1, …, *N* }. The relationships between the nodes of the graph is represented by the adjacency matrix **A** = [*A*_*ij*_] for binary graphs, and **W** = [*W*_*ij*_] for weighted graphs. In binary graphs, *A*_*ij*_ = 1 when nodes *i* and *j* are connected and *A*_*ij*_ = 0 when the nodes are not connected. For weighted graphs, *W*_*ij*_ represents the weight of the edge between nodes *i* and *j* and equals to zero when *i* = *j*. The degree matrix Δ is defined as the diagonal matrix with entries 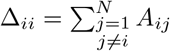, where Δ_*ii*_ is the degree of node *v*_*i*_. Similarly, the degree matrix Δ^*w*^ for weighted networks has diagonal entries 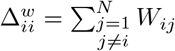.

For binary graphs the combinatorial Laplacian **L** is defined as **L** = Δ − **A**. The entries of **L** are given by

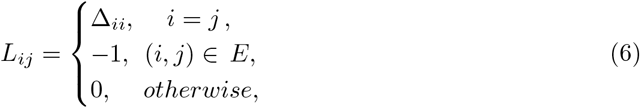

where Δ_*ii*_ is the degree of node *v*_*i*_. Similarly, for weighted graphs the Laplacian is defined as **L**^*w*^ = Δ^*w*^ − **W**.

### Graph Theoretic Measures

Complex networks can be characterized using graph theoretic metrics such as the clustering coefficient, characteristic path length, global efficiency, small world parameter and small world propensity [35, 36]. In this paper, we use graph theoretic measures defined for weighted networks as features for classification. Using graph theoretic measures defined for weighted networks circumvents the shortcomings associated with thresholding [4, 17, 18]. The features considered in this paper are as follows.

Clustering coefficient: The mean clustering coefficient is a measure of segregation and reflects mainly the fraction of clustered connectivity available around individual nodes. The clustering coefficient for a weighted network is defined as [37]:

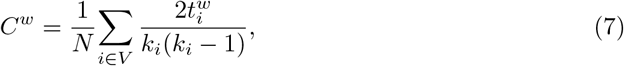

where 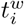 is the weighted geometric mean of the triangles around a node *i* defined as 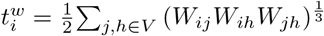 and *k*_*i*_ is the degree of node *i*.

Characteristic Path Length: The characteristic path length of the network is the average shortest path length between all pairs of nodes in the network. Path length in the brain network represents the potential routes of information flow between two different brain regions and quantifies the potential for functional integration [4]. For a weighted network, the characteristic path length is calculated as [4]:

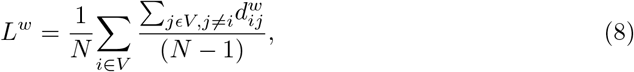

where 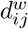 is the shortest weighted path length between node *i* and *j* defined as

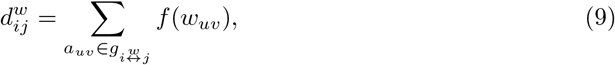

*f* refers to a map (e.g. an inverse function) from weight to length and 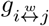 is the shortest weighted path between *i* and *j*.

Global Efficiency: The average inverse shortest path length is defined as the global efficiency of a network. It is a measure of functional integration similar to characteristic path length but can also be computed meaningfully for disconnected networks as an infinite path length results in zero efficiency [38]. The global efficiency for a weighed network is given by [38]:

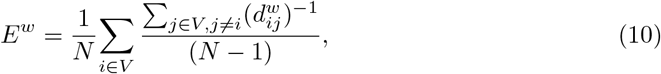

where 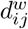 is the shortest weighted path length between node *i* and *j* defined by equation (9).

Small-World Parameter (SW): A network that has significantly more clusters than a random network but approximately the same characteristic path length as a random network is formally defined as a small-world network [39]. Small-world networks are simultaneously strongly clustered and integrated. This phenomenon of small worldness is captured by the small-world parameter which is the ratio of the normalized clustering coefficient to the normalized path length. For a weighted network, the small-world parameter is given as [4, 40]:

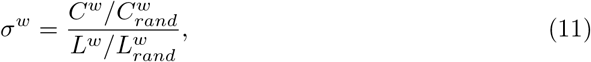

where *C* and *C*_*rand*_ are the clustering coefficients of the network and a random network with the same degree distribution, respectively, and *L* and *L*_*rand*_ are the characteristic path lengths of the network and a random network with the same degree distribution, respectively. The random networks are generated using the Erdos-Renyi model with the same number of nodes and connection density.

Small-World Propensity (SWP): Small world propensity is a measure that quantifies the level of small-worldness displayed by a network while accounting for the variation of network density [22]. SWP is measured by computing the deviation of the observed network’s clustering coefficient and characteristic path length from random (*C*_*rand*_, *L*_*rand*_) and lattice (*C*_*lat*_, *L*_*lat*_) networks designed with the same degree distribution and same number of nodes as follows:

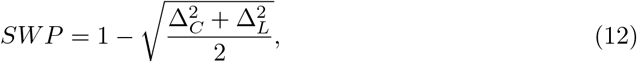

where 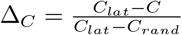, and 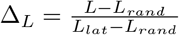.

## Methods

### Graph-to-Signal Transformation Based on the Resistance Distance Matrix

The objective of CMDS is to find a low-dimensional representation of the data such that the Euclidean distances between points are preserved [41]. In particular, for our application of transforming graphs into signals, the goal is to obtain coordinate vectors that preserve the defined distances on the graph [26].

In order to extract these coordinate vectors, first, the adjacency matrix **A** of a given network is transformed into a squared distance matrix, **D**^(2)^, which is consequently double centered as

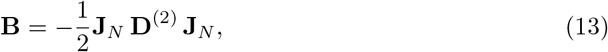

where **D**^(2)^ = **D** ◦ **D** is the entry-wise squared Euclidean distance matrix, 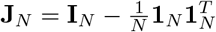 is a centering matrix, **I**_*N*_ is an *N* × *N* identity matrix, **1**_*N*_ is a *N* × 1 vector of ones, and *T* denotes the transpose. In order to preserve the positive definiteness of **B**, the matrix **D** needs to be a valid distance matrix and conditionally negative definite. In previous work [25, 42], CMDS has been limited to the transformation of binary networks where the distance **D** is based on the binary adjacency matrix **A**.

In order to extend the graph-to-signal transform based on CMDS to weighted graphs, we consider the resistance distance matrix of a graph, denoted as **R**. The resistance distance was introduced by Klein and Randic as an alternative to the shortest path distance for applications in chemistry [43]. It is inspired by basic circuit theory, where each edge on the graph represents a resistor with value 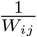 [44]. The resistance distance between nodes *i* and *j*, *R*_*ij*_, is defined for complete graphs through the Moore-Penrose pseudo inverse of the Laplacian, **L**, **L**^†^ [25], as

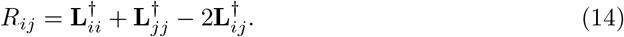

Each entry *R*_*ij*_ in **R** corresponds to the squared Euclidean distance between nodes *i* and *j* [45]. In a connected graph, *R*_*ij*_ ≤ *d*(*i, j*), where *d*(*i, j*) is the shortest path distance, and equality holds when there is only one path between *i* and *j* [46]. **R** is a valid squared Euclidean distance matrix as each entry *R*_*ij*_ satisfies the following rules [47]:

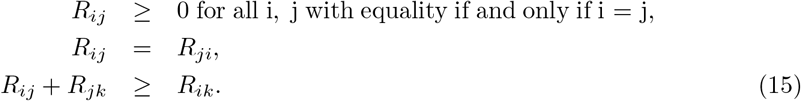

As a result, **R** can be directly substituted in (13) to obtain the corresponding **B** as

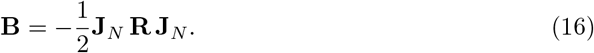

It can be shown that **B** is a positive semi-definite matrix with *rank*(**B**) = *C*, *C* ≤ *N*. Therefore, **B** has *C* nonzero eigenvalues, and *N* − *C* eigenvalues equal to zero.

The next step in graph-to-signal transformation is to perform the spectral factorization of **B**, resulting in 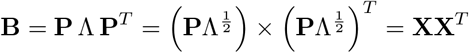, where Λ = *diag*(λ_1_, λ_2_, …, λ_*C*_) corresponds to the nonzero eigenvalues of **B**, with λ_1_ ≥ λ_2_ ≥ … ≥ λ_*C*_, **P** ∈ ℝ^*N* × *C*^, and **X** ∈ ℝ^*N* × *C*^. Based on **X**, a total of *C* signals of length *N* corresponding to the columns of **X** are obtained. The *i*th signal **x**_*i*_ ∈ ℝ^*N* × 1^ is defined as the *i*th column of **X** with *i* = 1, 2, …, *C*. In this paper, we will refer to **x**_*i*_*s* as the signals representing the network.

If the signals **X** are not distorted, it is possible to get back to the original network from the signals. First, the resistance distance matrix **R** can be recovered from the signals **x**_*i*_s through the computation of the squared Euclidean distance between the points as follows

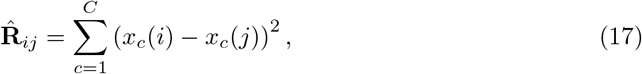

where 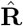 is the estimated **R**, *C* corresponds to the total number of components and *x*_*c*_(*i*) and *x*_*c*_(*j*) correspond to the *i*th and *j*th entries of the *c*th component. The original adjacency matrix can then be recovered from 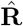, for both weighted and binary graphs following the procedure detailed in [48].

### Graph Signal Feature Extraction

In this section, we describe several well-known features adapted to graph signals. Along with common signal measures like Shannon Entropy (ShEn), skewness and kurtosis, we propose a new measure named graph spectral entropy (GSE) for quantifying the structural information of graphs based on the signals obtained from the networks. The extracted features are explained below.

Shannon Entropy (ShEn): Shannon entropy is a standard entropy measure widely used for signal analysis and it computes the order state of a signal by quantifying the probability density function of the distribution. Shannon entropy of the *i*th graph signal is computed as [49]:

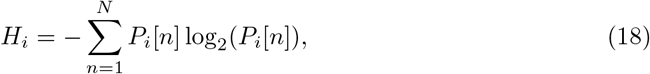

where *P*_*i*_ is the probability density function for the *i*th graph signal obtained through the histogram of the signal values. ShEn was computed for each of the graph signals, *x*_*i*_[*n*], *i* = 1, 2, …, *C*, generated through the graph-to-signal transformation. The average entropy, 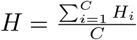 over all signals was extracted as a feature for consequent analysis. Skewness and Kurtosis: Skewness (S) [50] and Kurtosis (K) [51] of a signal are measures of the third and fourth moments, respectively and are defined as 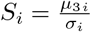 and 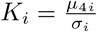, where *μ*_3*i*_ is the third central moment, *μ*_4*i*_ is the fourth central moment and *σ*_*i*_ is the standard deviation of the *i*th signal. The average skewness 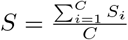 and average kurtosis 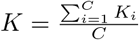 measured over all the signals were considered as two features of the graph signal.

Graph Spectral Entropy: We propose a new graph entropy measure based on the spectra of the graph signals. In particular, we propose to compute the graph entropy based on the normalized power spectrum of 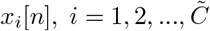, where we consider the 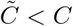 signals with highest energy. This parameter is selected empirically similar to the selection of the total number of factors in Principal Components Analysis (PCA). The magnitude spectrum of the *i*th signal is defined as 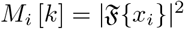, where 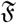 denotes the discrete Fourier transform, 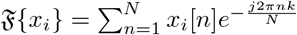. The normalized power spectrum of the *i*th signal for the positive frequencies is computed as 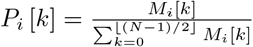, where *k* = 0, 1, …, ⌊(*N* − 1)/2⌋ corresponds to discrete frequency bins [52]. The normalized graph entropy for the *i*th graph signal is defined as

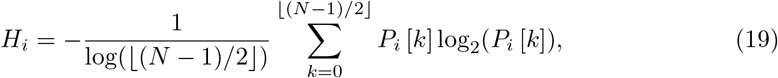

where 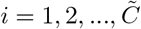 [52]. Since (19) refers to the Shannon entropy, it is bounded as 0 ≤ *H*_*i*_ ≤ log(*N*^2^/2). We propose to use the normalized power spectrum rather than the original signals for entropy computation since computing the Shannon entropy directly on the signals does not necessarily provide information about the network’s structural content. For example, for a structured network such as the ring network, the corresponding signals are pure sinusoids [25, 53], with almost uniform histograms resulting in high entropy. On the other hand, the power spectrum of a sine wave is well localized at a particular frequency thus its Shannon entropy is theoretically zero. This is consistent with the intuition that a ring network is deterministic and thus, should exhibit low entropy. Thus, the lower bound of *H*_*i*_ is achieved when the distribution is an impulse, and the upper bound occurs when the distribution is uniform. In terms of graph structures, the lower bound corresponds to the ring lattice and the upper bound corresponds to a random network.

In order to account for the variation in the network entropy as the probability of attachment varies, we propose to weigh the entropy of each graph signal using its energy using weights 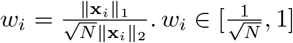, using the fact that 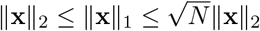. These weights are normalized across signals as 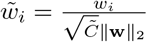, where 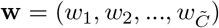. We define the weighted graph spectral entropy (GSE) as

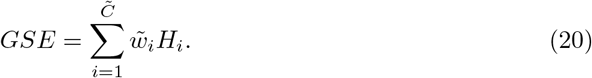

This definition of network entropy is independent of graph theoretic measures and the eigenspectrum of the adjacency matrix. The structural information of the network is thus obtained from the signals that already contain the network topological information.

## Results

### Simulations

#### Graph-to-signal transformation for binary networks

We first compare the proposed distance measure, **R**, to **D** for binary networks. For this purpose, we qualitatively compare the signals obtained from multiple binary networks. First, we simulate two ring networks with *N* = 128 nodes and average degrees *K* = 2 and *K* = 10. Fig. 1 (a) and (b) show the graph signals with the highest eigenvalue obtained from **R** and **D**, respectively. As expected, the signals based on the resistance distance matrix are sinusoidal signals (Fig. 1 (a)). From these figures, it is observed that the amplitude of signals obtained from **R** is inversely proportional to the average degree, *K*, yielding a higher amplitude when *K* = 2 and a smaller amplitude when *K* = 10. On the other hand, **D** cannot distinguish between ring networks with varying average degree.

We also compared both methods for an Erdős-Rènyi binary graph for a probability of attachment *p* equal to 0.5. For the original distance matrix **D**, the signals are random signals (Fig. 1 (d)), as previously shown in [27]. On the other hand, signals estimated from **R** still exhibit a random structure, with peaks that are inversely proportional to *p* (Fig. 1 (c)). The location of these peaks corresponds to the nodes with the smallest degree, i.e. the largest peak occurs in the first signal and corresponds to the node with the smallest degree. For the resistance distance, a node with small degree will have a high resistance distance between it and the remaining nodes in the network. Therefore, signals obtained from the transformation of binary networks through the resistance distance are more informative than those obtained from **D**.

**Fig 1.**
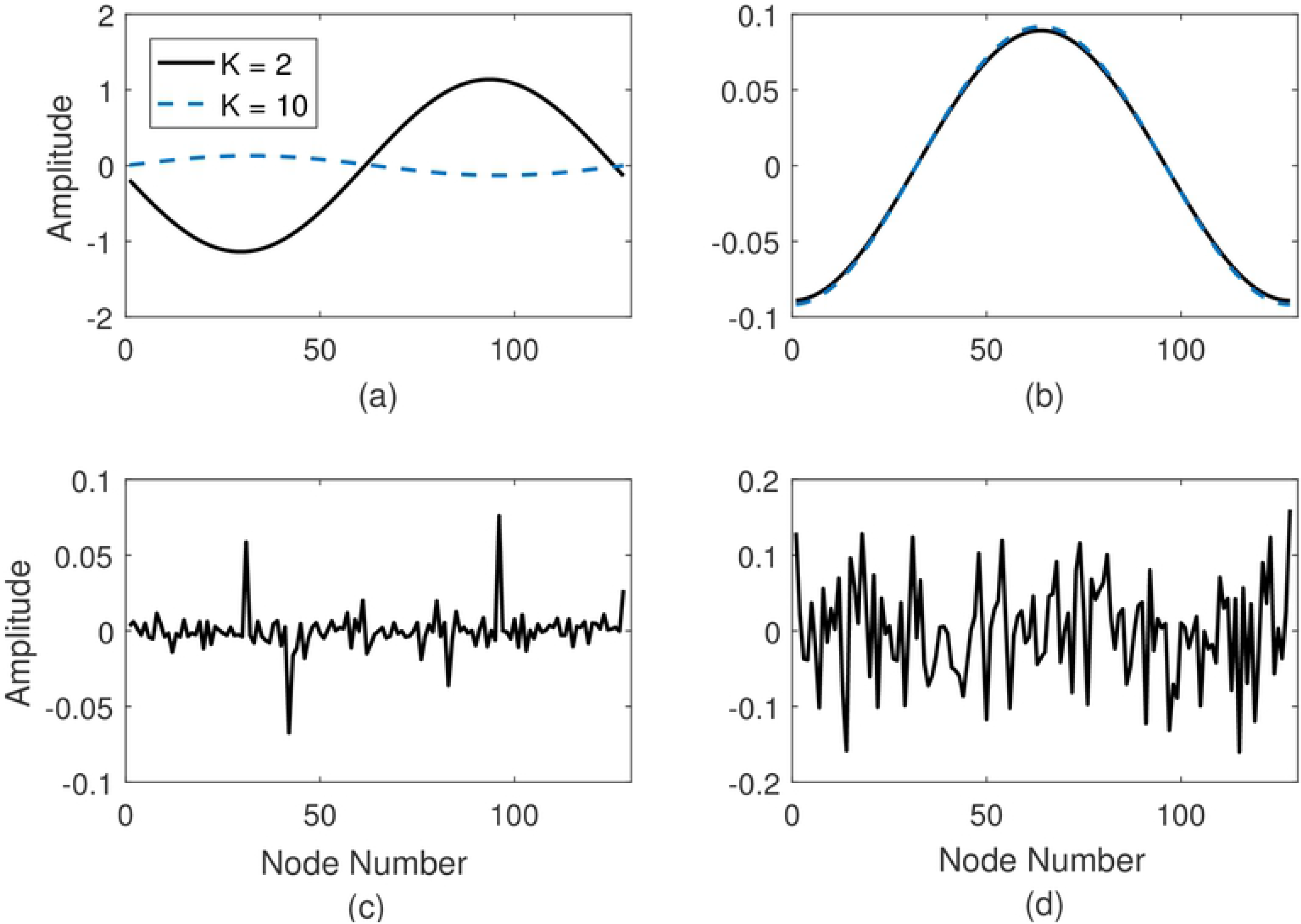
Graph signals from binary networks. Top: Graph-signals extracted from a ring lattice network with degree *K* = 2 and *K* = 10 using; A. Resistance distance, **R**, B. Distance **D**; Bottom: Graph-signals extracted from an Erdős-Rènyi network with probability of attachment *p* = 0.5 using; C. Resistance distance, **R**, D. Distance **D**. For all networks *N* = 128.

#### Graph-to-signal transformation for weighted graphs

The proposed transformation was also assessed on weighted networks. Fig. 2 (a) shows the signals resulting from a small-world network with average degree *K* = 6, and *N* = 128 nodes. As seen in Fig. 2 (a), for a network with a low rewiring probability, *p* = 0.1, the resulting signals are sinusoidal signals with some noise. This is consistent with previous work on binary networks [25], where it has been shown that the small-world network is equivalent to a ring network plus noise.

In addition to the small-world network, we investigated the graph-to-signal transformation of a weighted stochastic block network consisting of 200 nodes and with fixed probability of attachment, *p* = 0.3 and 3 clusters (Fig. 2 (b)). The weights are assigned randomly from the uniform distribution in the interval [0, 1]. It can be observed from these figures that the first *K* − 1 signals reveal the clusters’ structure, and the *K*th signal is an impulse. In addition, the size of each cluster can be inferred from the support of the constant regions in the first *K* − 1 signals. Thus, the proposed approach effectively transforms weighted networks into signals and reflects structural properties of the networks.

**Fig 2.**
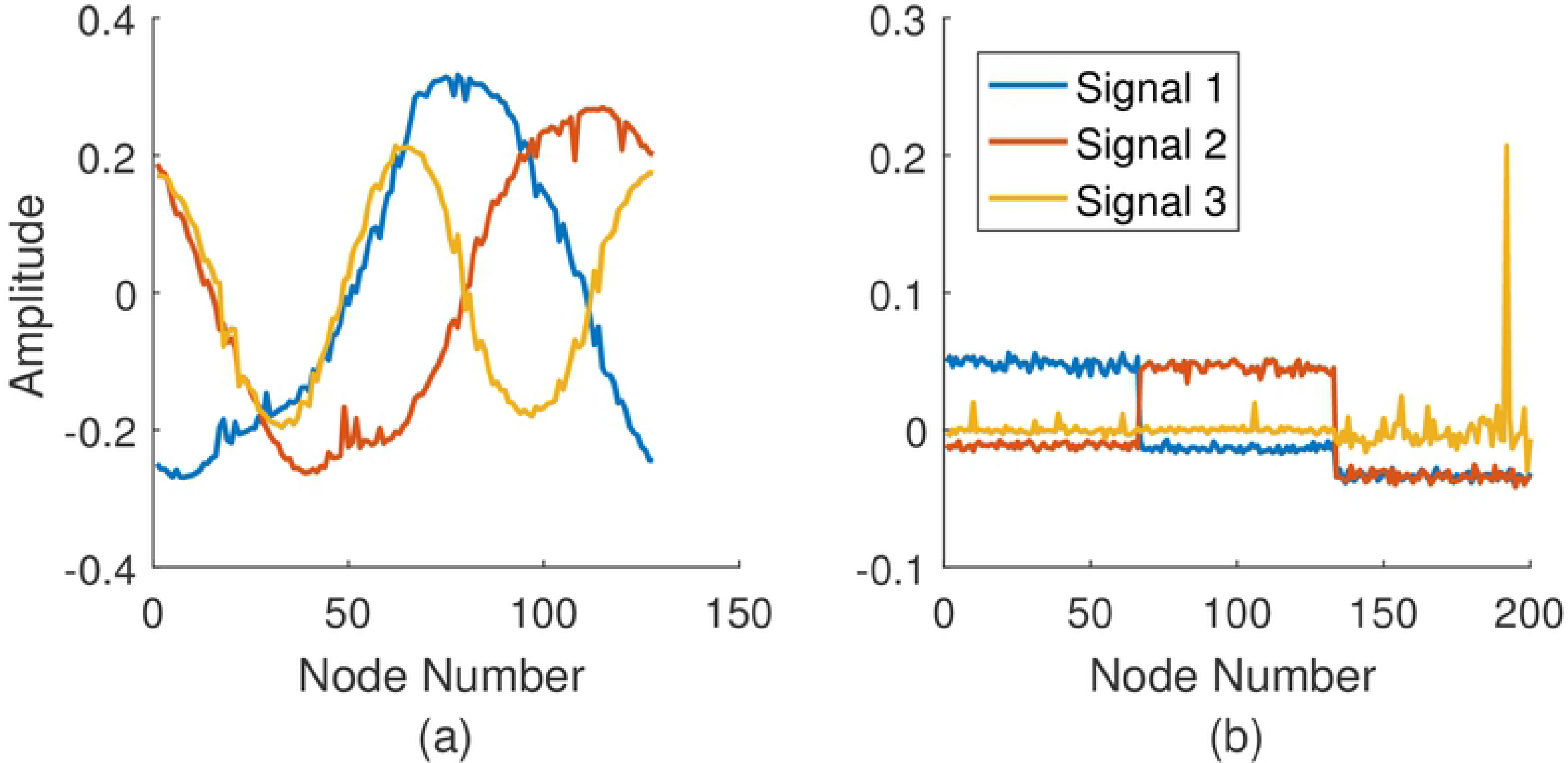
Graph signals from weighted networks. First three signals from A. A weighted small-world network with *K* = 6, *p* = 0.1 and *N* = 128 nodes; B. A weighted stochastic block network with probability of attachment *p* = 0.3 and *N* = 200 nodes.

### EEG data

In this paper, we analyze an EEG dataset from a previously published cognitive control-related error processing study [54]. The study was designed following the experimental protocol approved by the Institutional Review Board (IRB) of the Michigan State University. The data collection was performed in accordance with the guidelines and regulation established by this protocol. Written and informed consent was collected from each participant before data collection.

The experiment consisted of a speeded-reaction Flanker task [55] in which subjects identified the middle letter on a five-letter string, being congruent (e.g. MMMMM) or incongruent (e.g. MMNMM) with respect to the Flanker letters. Flanker letters (e.g. MM MM) were shown during the first 35 ms of each trial, and during the following 100 ms the Flanker and target letters were shown on the screen. This was followed by an inter-trial interval of variable duration ranging from 1200 ms to 1700 ms. A total of 6 blocks consisting of 80 trials composed the experiment, and letters were changed between blocks. EEG responses were recorded by the 64 electrodes ActiveTwo system (BioSemi, Amsterdam, The Netherlands). Trials containing artifacts were rejected and volume conduction was reduced through the Current Source Density (CSD) Toolbox [56]. A total of 18 subjects were considered for the analysis, for which the total number of error trials ranged from 20 to 61. The same number of correct responses was chosen randomly. The sampling frequency was 512 Hz. Fig. 3 shows the event-related potentials for error and correct responses, i.e. error-related negativity (ERN) and correct-related negativity (CRN), from electrode FCz averaged over trials and subjects. As can be seen from this figure, ERN has a larger negative amplitude with the peak within 0-100 ms, where 0 refers to the response time.

In this paper, we are interested in studying the differences in the FCNs corresponding to error-related negativity (ERN) and the correct-related negativity (CRN) through a classification task. The FCNs for both error and correct responses were constructed using the bivariate phase-locking value (PLV) between pairs of electrodes [57]. Previous studies have shown that the ERN is associated with increased synchronization in the theta band (4-8 Hz) between electrodes in the central and lateral frontal regions [54, 58, 59]. For this reason, a FCN was constructed for each subject by averaging the PLV over the time window 25-75 ms and the frequency bins corresponding to the theta band. A total of 58 signals are considered from the graph-to-signal transformation of each FCN.

**Fig 3.**
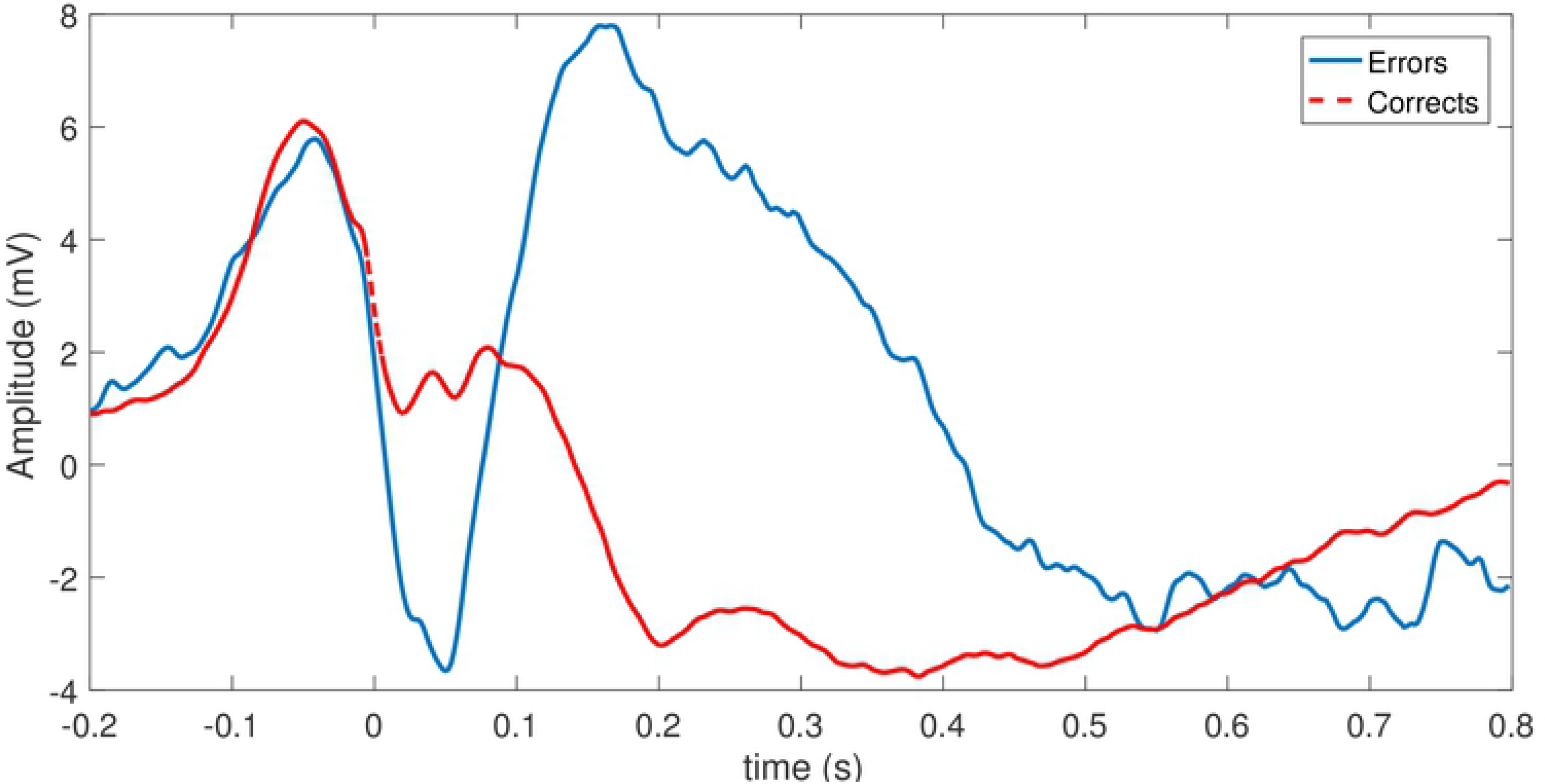
Average error and correct responses at FCz across all trials and all subjects.

#### Graph-to-Signal Transformation of FCNs

The average FCNs constructed from ERN and CRN waveforms for the 25-75 ms time interval and the theta frequency band (4-8 Hz) were transformed into signals using (16). For illustration purposes, we show the first six graph signals corresponding to the correct and the error responses in Figs. 4 (a) and Fig. 4 (b), respectively. We focus on the first six signals obtained from this transformation as the eigenvalues of the matrix **B** in (16) drop off significantly after the sixth eigenvalue. As the graph signals are a function of the nodes or different electrodes, the location of the peaks of the graph signals signify the distribution of spatial activity. It can be observed from Fig. 4 that while the energy of the graph signals from CRN is distributed uniformly across the 58 brain regions, the energies of the ERN graph signals are more concentrated within the first 20 electrodes, which correspond to the frontal and frontal-central regions. This implies that right after an error response most of the brain activity centralizes within the frontal regions. This is line with prior work indicating the role of prefrontal cortex during ERN [59, 60].

**Fig 4.**
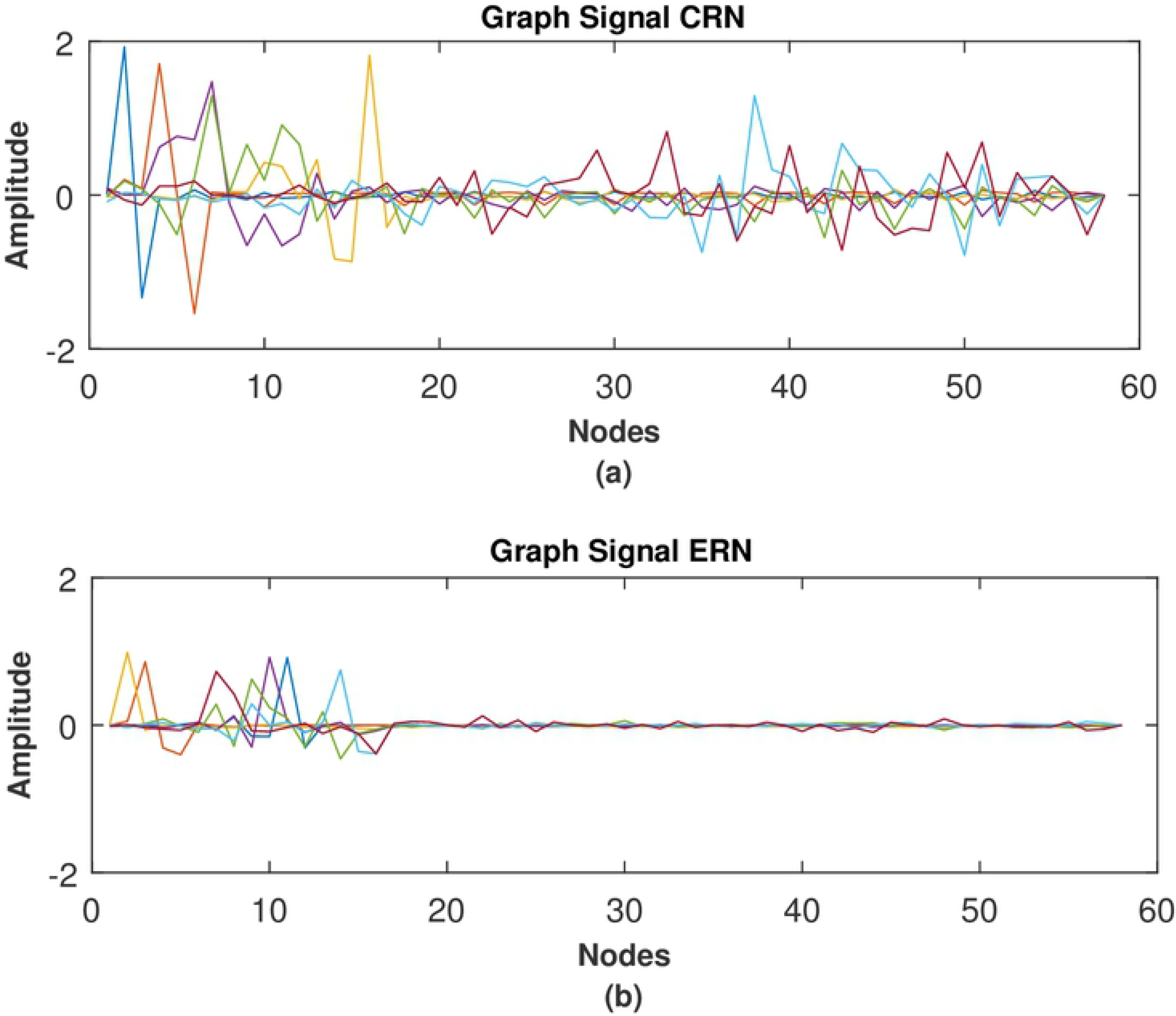
Graph Signal representation. The first six signals obtained from graph-to-signal transformation of A. CRN networks; B. ERN networks.

Fig. 5 (a) and Fig. 5 (b) show the magnitude spectra of the signals corresponding to average error and correct FCNs. For the average FCN constructed from error responses, the frequency content of the signals increases with the signal number, suggesting an organized structure such as ring networks. On the other hand, the spectra of graph signals corresponding to correct responses suggest a random network structure.

**Fig 5.**
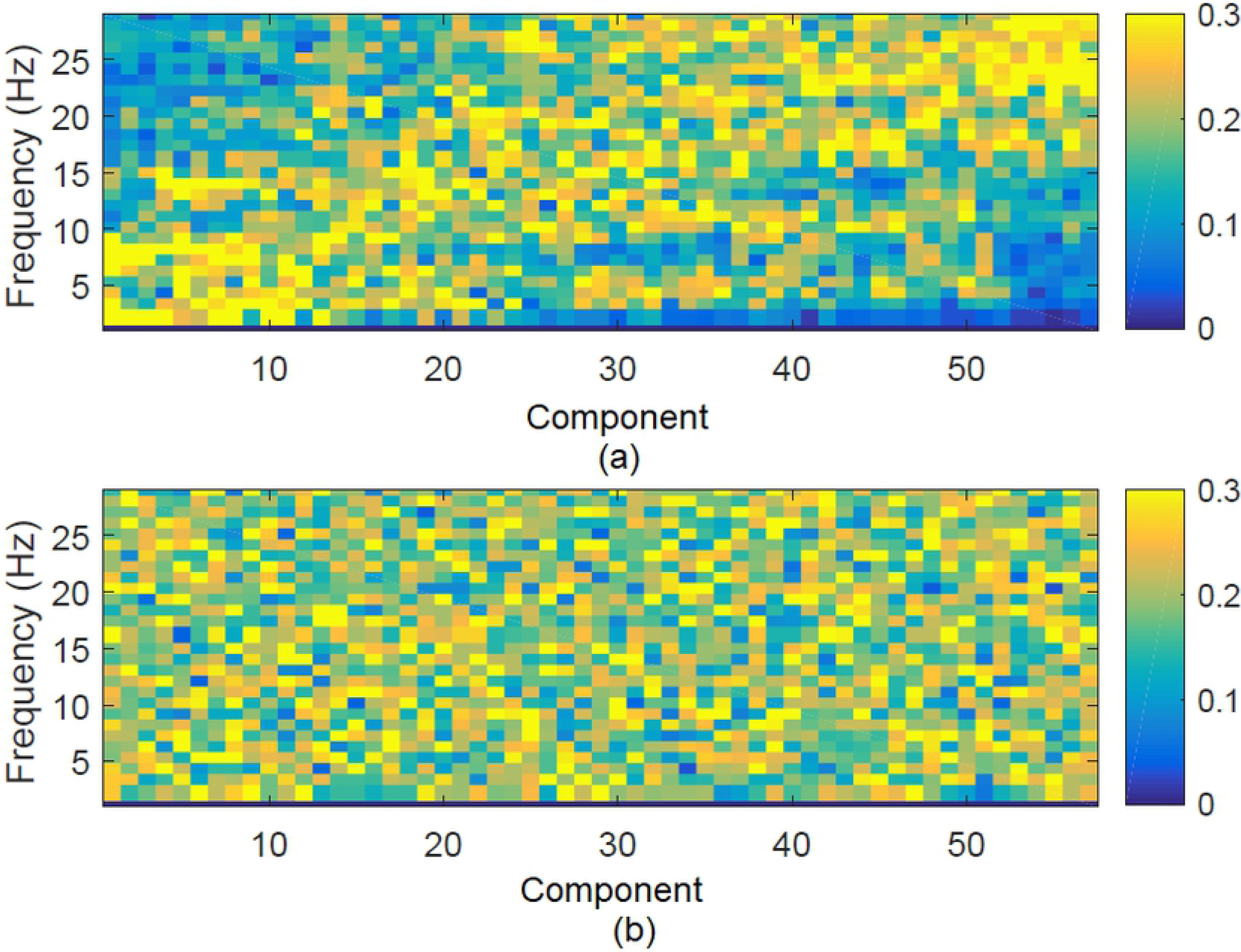
Magnitude Spectrum Representation. Magnitude Spectrum for each signal obtained through graph-to-signal transformation for A. Error responses; B. Correct responses.

#### Classification of FCNs

In this section, we evaluate the classification power of the features extracted from graph signals and compare these features with conventional graph theoretic measures. The comparison was conducted by performing a classification task between ERN and CRN networks based on the features extracted from these networks through graph theoretic measure and graph-signal transform. For a comprehensive comparison, we employed a set of classifiers including support vector machines (SVM), linear discriminant analysis (LDA), logistic regression and k-nearest neighbor (kNN) (with *k* = 20). The accuracy of each classification method was determined based on its prediction accuracy on 5-fold cross-validation. Since the operation involves binary classification, sensitivity and specificity defined as follows were used as performance measures in addition to accuracy.

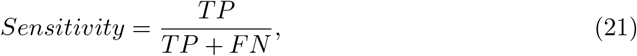

where *TP* is True Positive, *FN* is False Negative; and

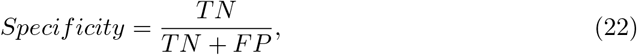

where *TN* is True Negative and *FP* is False Positive.

Table 1 shows the classification performance for graph theoretic features including clustering coefficient, path length, global efficiency, small world and small world propensity in terms of accuracy, sensitivity, and specificity for different classifiers. From these results, it can be seen that the small world propensity is the most effective graph theoretic feature. We also evaluated the accuracy of using all of the features together. An overall accuracy of 94.4% was obtained for linear SVM using all features. Moreover, the FCNs constructed from error responses exhibited significantly increased small-world (*p* = 0.00203, Wilcoxon rank-sum test) and small-world propensity measure (*p* = 0.0008280, Wilcoxon rank-sum test) compared to the FCNs from correct responses. This finding of decreased small-world characteristics for correct response networks is indicative of increased randomness and is in line with previous studies that reported increased small-worldness for ERN compared to CRN [21].

**Table 1.** Classification of ERN and CRN Functional Connectivity Networks using Graph Theoretic Features.

| Features | Linear SVM | LDA | Logistic Regression | KNN |
| --- | --- | --- | --- | --- |
| CC | 58.30% (Se:61%, Sp:56%) | 44.40% (Se:44%, Sp:44%) | 52.80% (Se:50%, Sp:56%) | 47.30% (Se:33%, Sp:61%) |
| PL | 61.10% (Se:56%, Sp:67%) | 52.80% (Se:50%, Sp:56%) | 55.60% (Se:44%, Sp:67%) | 38.90% (Se:44%, Sp:33%) |
| GE | 55.60% (Se:47%, Sp:64%) | 52.80% (Se:39%, Sp:67%) | 50.10% (Se:44%, Sp:56%) | 44.40% (Se:61%, Sp:28%) |
| SW | 81.70% (Se:84%, Sp:79%) | 78.90% (Se:79%, Sp:79%) | 76.10% (Se:79%, Sp:73%) | 81.70% (Se:96%, Sp:85%) |
| SWP | 84.20% (Se:88%, Sp:81%) | 80.10% (Se:82%, Sp:78%) | 77.20% (Se:80%, Sp:74%) | 80.40% (Se:86%, Sp:75%) |
| All | **94.40% (Se:100%, Sp:89%)** | **91.70% (Se:89%, Sp:94%)** | **88.90% (Se:83%, Sp:94%)** | **94.10% (Se:99%, Sp:89%)** |

Se: Sensitivity; Sp: Specificity; CC: Clustering Coefficient; PL: Characteristic Path Length; GE: Global Efficiency; SW: Small World Parameter; SWP: Small World Propensity.

Table 2 shows the classification performance for graph signal features including graph spectral entropy, Shannon entropy, skewness and kurtosis in terms of accuracy, sensitivity, and specificity for different classifiers. From these results, it can be seen that the graph spectral entropy was the most effective graph signal feature. An overall accuracy of 97.2% was obtained for linear SVM using all features for discriminating between ERN and CRN connectivity networks. It is an overall 3% improvement of accuracy compared to the graph theoretic measures. Moreover, FCNs from correct responses show higher entropy than FCNs from error responses and this difference is significant (*p* = 0.0000554, Wilcoxon rank-sum test). This is consistent with the fact that the error-related negativity is associated with increased synchronization which results in less random networks and hence lower network entropy.

**Table 2.** Classification of ERN and CRN Functional Connectivity Networks using Graph Signal Features.

| Features | Linear SVM | LDA | Logistic Regression | KNN |
| --- | --- | --- | --- | --- |
| GSE | 85.90% (Se:88%, Sp:83%) | 71.40% (Se:69%, Sp:72%) | 81.50% (Se:80%, Sp:83%) | 76.10% (Se:80%, Sp:72%) |
| ShE | 66.70% (Se:89%, Sp:56%) | 63.90% (Se:85%, Sp:43%) | 77.80% (Se:89%, Sp:67%) | 61.10% (Se:61%, Sp:61%) |
| S | 77.20% (Se:80%, Sp:74%) | 71.70% (Se:83%, Sp:68%) | 74.50% (Se:72%, Sp:77%) | 44.40% (Se:56%, Sp:33%) |
| K | 80.60% (Se:78%, Sp:83%) | 80.60% (Se:94%, Sp:67%) | 77.80% (Se:72%, Sp:83%) | 75.00% (Se:78%, Sp:72%) |
| All | **97.20% (Se:100%, Sp:94%)** | **97.20% (Se:100%, Sp:94%)** | **94.50% (Se:100%, Sp:90%)** | **94.20% (Se:94%, Sp:94%)** |

Se: Sensitivity; Sp: Specificity; GSE: Graph Spectral Entropy; ShE: Shannon Entropy; S: Skewness; K: Kurtosis.

Comparing Tables 1 and 2, it can be seen that three measures, small world parameter, small world propensity and graph spectral entropy, were the most effective features to differentiate between ERN and CRN networks. In order to determine which measure, as a continuous test statistics, best discriminates between error and correct networks, we have computed receiver operating characteristic (ROC) curve for each measure. ROC curve illustrates the performance of a binary decision boundary with the variation of the discrimination threshold. For each ROC curve, the area under the curve (AUC) was computed as it serves as a quantitative measure of the discrimination power of the test statistics. As shown in Fig. 6, graph spectral entropy has the highest AUC, indicating that among these three measures, graph spectral entropy is the most effective test statistic to discriminate between the two response types.

**Fig 6.**
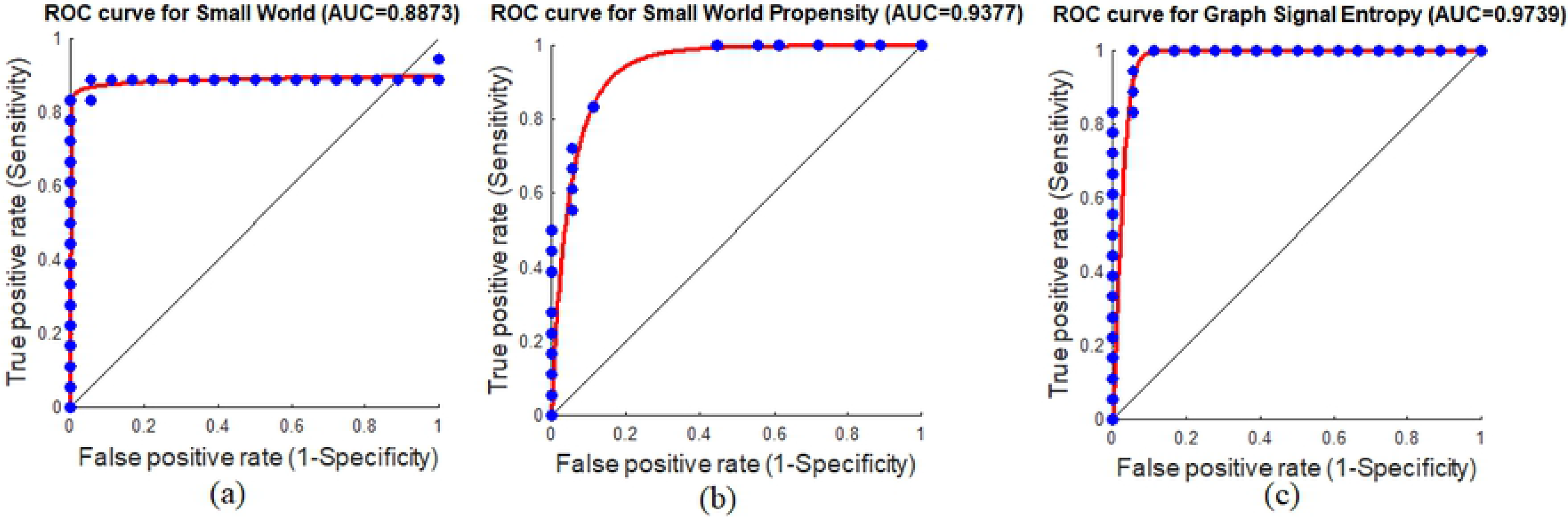
Area under the curve (AUC) and the ROC curves. A. Small-World; B. Small-World Propensity and C. Graph Signal Entropy measures.

## Conclusion

In this paper, we introduced a new graph-to-signal transformation for weighted FCNs. The signals obtained from this transformation were used to characterize the networks and to extract discriminative features. Results indicate that the features extracted from graph signals are more discriminative compared to conventional graph theoretic measures for classifying between FCNs corresponding to error and correct responses. In particular, the graph spectral entropy decreases during the ERN interval, while the entropy increases after correct responses. This indicates that ERN has a more modular structure implying increased segregation. This finding is in line with previous research showing more localized activity during ERN compared to CRN [61].

## Acknowledgments

The authors want to thank Dr. Jason S. Moser from the Department of Psychology at Michigan State University for providing the EEG data.

